# rawMSA: End-to-end Deep Learning Makes Protein Sequence Profiles and Feature Extraction obsolete

**DOI:** 10.1101/394437

**Authors:** Claudio Mirabello, Björn Wallner

## Abstract

In the last few decades, huge efforts have been made in the bioinformatics community to develop machine learning-based methods for the prediction of structural features of proteins in the hope of answering fundamental questions about the way proteins function and about their involvement in several illnesses. The recent advent of Deep Learning has renewed the interest in neural networks, with dozens of methods being developed in the hope of taking advantage of these new architectures. On the other hand, most methods are still based on heavy pre-processing of the input data, as well as the extraction and integration of multiple hand-picked, manually designed features. Since Multiple Sequence Alignments (MSA) are almost always the main source of information in *de novo* prediction methods, it should be possible to develop Deep Networks to automatically refine the data and extract useful features from it. In this work, we propose a new paradigm for the prediction of protein structural features called rawMSA. The core idea behind rawMSA is borrowed from the field of natural language processing to map amino acid sequences into an adaptively learned continuous space. This allows the whole MSA to be input into a Deep Network, thus rendering sequence profiles and other pre-calculated features obsolete. We developed rawMSA in three different flavors to predict secondary structure, relative solvent accessibility and inter-residue contact maps. We have rigorously trained and benchmarked rawMSA on a large set of proteins and have determined that it outperforms classical methods based on position-specific scoring matrices (PSSM) when predicting secondary structure and solvent accessibility, while performing on a par with the top ranked CASP12 methods in the inter-residue contact map prediction category. We believe that rawMSA represents a promising, more powerful approach to protein structure prediction that could replace older methods based on protein profiles in the coming years.

**Availability:** datasets, dataset generation code, evaluation code and models are available at: https://bitbucket.org/clami66/rawmsa

## 1 Introduction

Predicting the 3D-structure of a protein from its amino acid sequence has been one of the main objectives of bioinformatics for decades now [1], yet a definite solution has not been found yet.

The most reliable approaches currently involve *homology modeling*, which allows a known protein structure to be assigned an unknown protein provided that there is a detectable sequence similarity between the two. When homology modeling is not viable, *de novo* techniques are needed, either based on physical-based potentials [2] or knowledge-based potentials [3–6].

In the first case, an energy function is used to approximate the amount of free energy in a given protein conformation along with a search function that tries different structural conformations in order to minimize such energy function [7]. Unfortunately, proteins are very large molecules, and the huge amount of available conformations, even for relatively small proteins, makes it prohibitively expensive to fold them even on customized computer hardware [8].

Knowledge-based potentials, on the other hand, can be learned from statistics or machine learning methods to infer useful information from known examples of protein structures. This information can be used to constrain the problem, thus greatly reducing the number of samples that need to be evaluated when dealing exclusively with physics-based potentials.

In the last couple of decades, a variety of machine learning methods have been developed to predict a number of structural properties of proteins: secondary structure (SS) [9–14], relative solvent accessibility (RSA) [14–17], backbone dihedrals [18], disorder [14, 19, 20], disorder-to-order transition [21, 22],contact maps [23–26], and model quality [27–30].

The most important information used by most (if not all) methods above is a multiple sequence alignment (MSA) for homologous sequences to the target protein. The MSA consists of aligned sequences and to allow for comparisons and analysis of MSAs, they are compressed into position-specific scoring matrices (PSSM), also called sequence profiles, using the fraction of occurrences of different amino acids in the alignment for each position in the sequence. The sequence profile describes the available evolutionary information of the target protein and is better than a single sequence representation, often allowing for a significant increase in the prediction accuracy [31, 32].

An obvious limitation of compressing an MSA into a PSSM is the loss of information that could be useful to obtain better predictions. Another potential issue is that whenever the MSA contains few sequences, the statistics encoded in the PSSM will not be as reliable and the prediction system may not be able to distinguish between a reliable and an unreliable PSSM.

SS, RSA and similar properties are sometimes used as intermediate features to constrain and help the prediction of more complex properties in a number of methods [33–35]. An example of this comes from the methods used for the prediction of inter-residue contact maps, where evolutionary profiles are integrated with predicted SS and RSA to improve performance [16, 36, 37].

More recently, contact map prediction methods have been at the center of renewed interest after the development of a number of techniques to analyze MSAs in search of direct evolutionary couplings. These methods have led to a big leap in the state of the art [38–40]. However, their impressive performance is correlated with the number of sequences in the MSA, and is not as reliable when few sequences are related to the target. This means that evolutionary coupling methods have not completely replaced older machine learning-based systems, but have been integrated, usually in the form of extra inputs, along with the previously mentioned sequence profiles, SS and RSA, into even more complex machine learning systems. At the same time, the Deep Learning revolution has proved to be a useful tool for better integrating the growing number and complexity of input features [41–43].

However, one might argue that this kind of integrative approach, combining individually derived features, ignores a key aspect of deep learning, i.e. that features should be automatically extracted by the network rather than being provided to the network as inputs [44]. If we wanted to take advantage of deep learning by using it in the same way it is employed for tasks such as image classification, one idea could be to provide a raw MSA input, which is the basic, lowest level input that most methods use, and let the deep network take care of the feature extraction, avoiding the loss of potentially useful information through the compression into profiles. On the other hand, a MSA is not an image or an audio track, and there is no native way of feeding such a large block of strings as input to a deep network.

In this work we try to overcome this hurdle and introduce a new system for the *de novo* prediction of structural properties of proteins called rawMSA. The core idea behind rawMSA is to borrow from the field of natural language processing a technique called *embedding* [45], which we use to convert each residue character in the MSA into a floating-point vector of variable size. This way of representing residues is adaptively learned by the network based on *context*, i.e. the structural property that we are trying to predict. To showcase the idea, we designed and tested several deep neural networks based on this concept to predict SS, RSA, and Residue-Residue Contact Maps (CMAP). Our test results show that rawMSA matches or outperforms the state of the art in all three applications.

## 2 Methods

### 2.1 Inputs

Unlike the classical machine learning methods for the prediction of protein features, rawMSA does not compress the Multiple Sequence Alignment into a profile but, rather, uses the raw aligned sequences as input and devolves the task of extracting useful features to the deep network. The input to the deep network is a flat FASTA alignment file. Before it is passed to the input layer of the neural network, each letter in the input file (including gaps) is mapped to an integer ranging from 1 to 25 (21 standard residues plus the non-standard residues *B*, *Z*, *X* and the gap placeholder, –). If the alignment file of a protein of length *L* contains *N* sequences (including the target, or “master” sequence), it is translated to an array of *L × N* integers. The master sequence occupies the first row of the array, while the following rows contain all the aligned sequences, in the order of output determined by the alignment software. Since the MSAs can become quite “deep” when aligning larger protein families (up to tens of thousands of sequences), a threshold is set so that no more than *Y* sequences are used, while the rest (usually alignments at higher E-values) are discarded. For details on the depth threshold, see the “Architecture” paragraph.

When training on or predicting SS or RSA, a sliding window of width 31 is applied to the MSA so that *L* separate windows of size 31 × *Y*, one for each residue in the master sequence, are passed to the network. The central column in the window is occupied by the residue in the master sequence for which a prediction is being made and the corresponding aligned residues from the other sequences. Zero-padding at the N- or C- terminals of the master sequence is applied when the number *N* of aligned sequences is such that *N < Y* or if the master sequence’s length *L* is such that *L < X* so that the window is of the correct shape.

### 2.2 Architecture

We developed two different architectures for three different applications “SS-RSA” for the SS and RSA prediction and “CMAP” for the contact map prediction In Fig. 1 we show an example of the network architecture. The networks trained in this work might use different numbers of convolutional, fully connected or BRNN layers, as well as slightly different parameters, but they all share this same basic structure.

**Fig 1.**
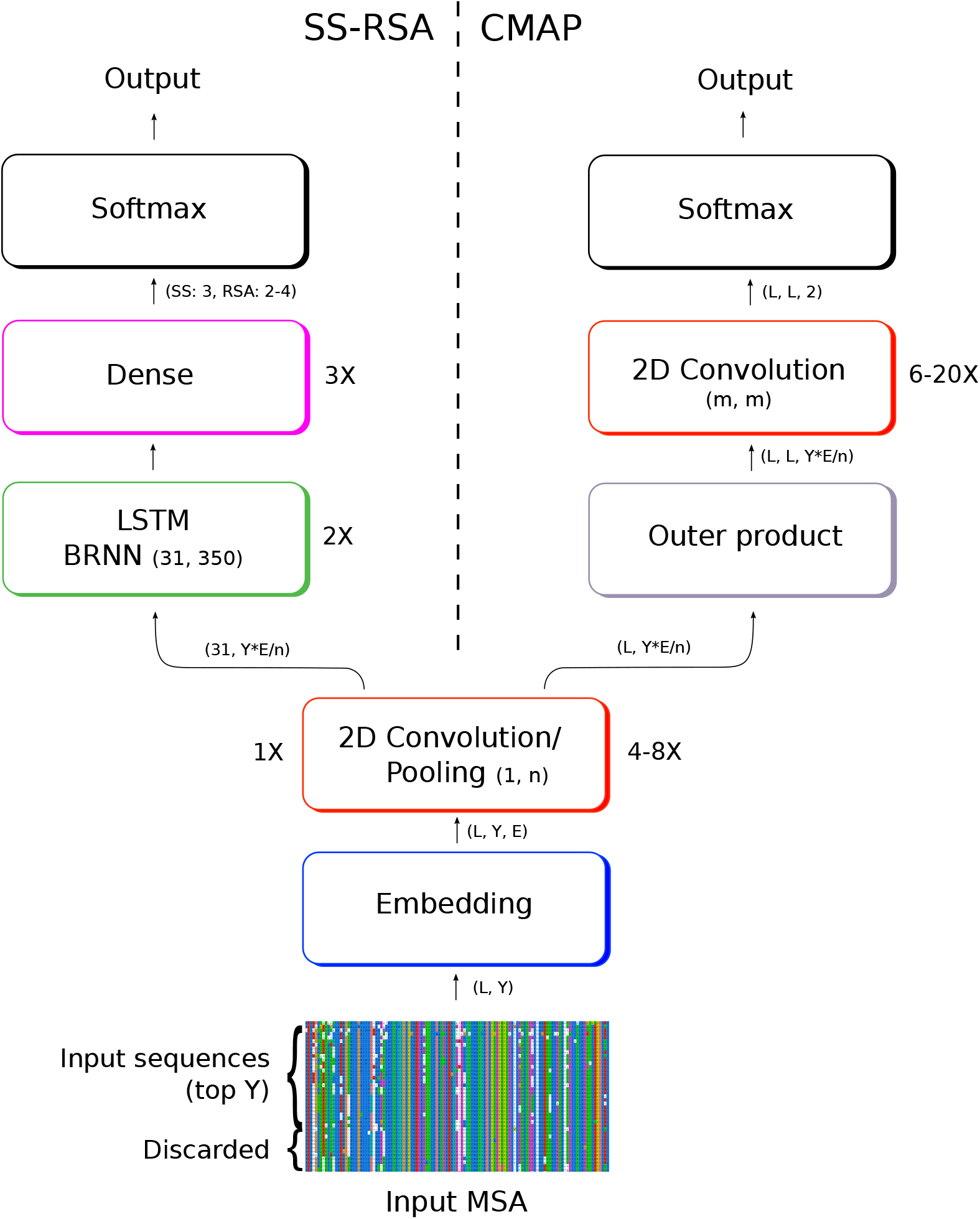
Schematics of two networks developed on rawMSA. On the left, the “SS-RSA” network predicts the secondary structure and relative solvent accessibility of each amino acid; on the right the “CMAP” network predicts the full contact map of the protein. The first layers are in common between the SS-RSA and CMAP architectures, although with slightly different settings, and provide the basis for the rawMSA approach.

Since with rawMSA we abandoned the use of sequence profiles, which are also useful to represent the amino acid information in a computer-friendly format, i.e. a matrix of floating points, we needed to come up with a way of representing the input. In this case, where the inputs can be very large (up to hundreds of thousands of amino acids), categorical data cannot be translated with sparse, memory inefficient techniques such as *one-hot encoding*.

To resolve this issue, the first layer of rawMSA is a trained, shallow two-layer neural network called an *embedding layer*. Embeddings are a compact way of representing a discrete set of inputs into a continuous space [45]. This technique is widely used in natural language processing where the inputs are made of a sequence of words taken from a dictionary and mapped to an n-dimensional space in a vector of floats of size *n*. When dealing with word embeddings in natural language, words that represent similar concepts, at least in a certain context, will be in close proximity in the output space. A similar idea might also be very useful when dealing with the discrete set of amino acids [46], since they also share context-dependent similarities. For example, when it comes to the context of secondary structure, if we look at the Chou-Fasman amino acid propensities table [47], glutamic acid and methionine are both strongly associated with alpha-helices, so it might be useful that such amino acids are represented by similar vectors when predicting secondary structure.

The embedding layers in rawMSA output a vector of size 10 to 30, depending on the model, for each input residue in the alignment. The embedding layer is used both in the SS-RSA and the CMAP networks.

#### 2.2.1 SS-RSA

In the case of SS-RSA, a 2D convolutional layer is stacked on top of the embedding layer, followed by a max pooling layer. The convolutional layer has a number of filters equal to the dimensionality of the embedding space. The convolution filters have the shape of column vectors, rather than square matrices as is usually the case, so the size of the convolution windows varies between 1 × 10 to 1 × 30 depending on the model. This means that convolution is performed along each column in the MSA and the information does not spread across columns (i.e. across adjacent residues in the input sequence). Pooling is performed selecting the maximum value in a window of the same size. In this way, if the dimension of the input is 31 residues by 500 alignments before embedding and 31 × 500 × 10 after embedding, this is reduced to a vector of size 31 × 50 × 10 following the convolution and pooling layers, if the convolution and pooling windows are of size 1 × 10. The convolutional and pooling layers are followed by a stack of two Long Short-Term Memory (LSTM) bidirectional recurrent layers, where each LSTM module contains 350 hidden units. The final three layers are fully connected, with softmax layers to output the classification prediction for SS (3 classes: Helix, Strand, Coil) or RSA (4 or 2 classes), depending on the model. Dropout is applied after each recurrent or dense layer to avoid overfitting, with variable fractions of neurons being dropped depending on the model (0.4 to 0.5). All the convolutional layers have ReLU activations and the outputs are zero-padded to match the two first dimensions of the inputs.

#### 2.2.2 CMAP

For CMAP, the network predicts a whole contact map of size *L × L* for a protein of length *L*. The input, in this case, is not split in windows, but we use the whole width MSA at once, while the depth is cut at the *Y* top alignments. The network for the first part of CMAP is similar to SS-RSA, with an embedding layer followed by up to six (rather than only one) 2D convolution/max pooling layers along each MSA column. In this case though, the output of the network is a contact map with shape *L × L*, so the preceding layers should represent the interaction between pairs of residues. This change of dimensionality is performed with a custom layer that performs a high-dimensional outer product of the output by the first stack of convolutional layers (*H*) with itself (*H* ⊗ *H*). This operation generates a four-dimensional hidden tensor of shape *L × L × F × S*, where *F* and *S* are the last two dimensions of the hidden tensor before the outer product. This output is then reshaped to a three-dimensional tensor of shape *L × L × (F*S*) and passed to a new stack of six to 20 (depending on the model) 2D convolutional layers with squared convolutions of varied size (3x3, 5x5, 10x10) and number of filters (10 to 50). The last convolutional layer has shape *L × L ×* 2 and is followed by a softmax activation layer that output the contact prediction with a probability from 0 to 1. All the convolutional layers have ReLU activations and the outputs are zero-padded to match the two first dimensions of the inputs. Batch normalization is performed in the outputs of the convolutional layers in the CMAP network.

### 2.3 Training

We wrote the code for rawMSA in Python using the Keras library [48] with TensorFlow backend [49]. Training and testing were performed on computers equipped with NVIDIA GeForce 1080Ti, TESLA K80, and Quadro P6000 GPUs.

The training procedure was run including one protein in each batch block, regardless of the size, and using an RMSprop optimizer with sparse categorical cross-entropy as loss function. The SS-RSA network was trained for five epochs, while the CMAP network was trained for up to 200 epochs. During training, a random 10% of the training samples were reserved for validation, while the rest were used for training. After training, the model with the highest accuracy on the validation data was used for testing.

### 2.4 Data sets

The data set is composed of protein chains extracted from a 70% redundancy-reduced version of PDB compiled by PISCES [50] in April 2017 with minimum resolution of 3.0 Å and R-factor 1.0. This set contains 29,653 protein chains.

#### 2.4.1 Avoiding homolog contamination

When training our networks, we want to make sure that the testing and training sets are rigorously separated so that no protein in the test set is too similar to any protein that the network has already “seen” during the training phase.

While most secondary structure and solvent accessibility prediction methods have been using 25-30% sequence identity as the threshold to separate testing and training sets [51–54], this practice has been discouraged as it has been shown that it is not sufficient to avoid information leakage [55]. This is apparently also valid for raw MSA inputs, as in our tests separating sets at 25% sequence identity yields higher accuracies compared to our final results (data not shown).

To correctly split training and testing sets, we used two databases based on a structural classification of the proteins: ECOD [56] and SCOPe [57] databases to assign one or more super families to each of the protein chains in the initial set. Then, we removed any chains that were related to more than one superfamily. The set generated from ECOD contains 16675 proteins (ECOD set), while the one generated from SCOPe contains 9885 proteins (SCOPe set). The SCOPe set contains fewer proteins than the ECOD set since SCOPe has a lower coverage of the PDB. We split each set into five subsets by making sure that no two proteins from the same superfamily were placed in two separate subsets. This ensured that the respective MSA inputs would not be too similar to each other and is the recommended practice when training neural networks using sequence profiles, which are themselves extracted from MSAs, as inputs [55].

We used the SCOPe subsets to perform five-fold cross-validation on SS-RSA. We also used one of the ECOD subsets to train and validate CMAP, where the validation set was used to determine when to stop training to avoid overfitting, and to select the models that would be ensembled and tested (i.e. the models with the lowest validation error).

#### 2.4.2 Multiple Sequence Alignments

The MSAs for both SS-RSA and CMAP were obtained with HHblits [58] by searching with the master sequence against the HMM database clustered at 20% sequence identity from February, 2016 for three iterations, with 50% minimum coverage, 99% sequence similarity threshold, and 0.001 maximum E-value.

We also obtained a second set of MSAs by running JackHMMER [59], for three iterations and 1*e −* 3 maximum E-value on the UniRef100 database from February 2016.

The HHblits alignments were used to train and test the SS-RSA networks. The HHblits alignments were used to train the CMAP network, while both the HHblits and the JackHMMER alignments were used as inputs when ensembling CMAP networks (see Ensembling section below), as it improved the prediction accuracy. This approach was also tried for the SS-RSA network, but no improvement was observed.

#### 2.4.3 Test sets

We labeled the CMAP data sets by assigning a native contact map to each protein. A contact was assigned to a couple of residues in a protein if the Euclidean distance between their *Cβ* atoms (*Cα* for Gly) in the crystal structure from the PDB was lower than 8 Å. Otherwise, the two residues were assigned the non-contact label.

We tested CMAP on the last CASP12 RR (Residue-Residue) benchmark [60], which is composed of 37 protein chains/domains of the Free Modeling class, i.e. protein targets for which no obvious protein homologs could be found at the time of the experiment (May-August 2016). To ensure a fair comparison with the predictors which participated in CASP12, we performed the benchmark in the same conditions to which all the other predictors where subjected at the time of the CASP experiment. We made sure that all protein structures (from Apr, 2016) in the training set (Apr, 2016) and sequences (from Feb, 2016) in the HHblist HMM and UniRef100 databases were released before CASP12 started.

To test SS-RSA, we calculated the secondary structure (SS) and the Relative accessible Surface Area (RSA) with DSSP 2.0.4 [61]. We reduced the eight SS classes (G, H, I, E, B, S, T, C) to the more common three classes: Coil, Helix, Extended (C, H, E). We used the theoretical Maximum Accessibility Surface Area (Max ASA) defined in [62] to calculate the RSA from the absolute surface areas (ASA) in the DSSP output and we used [0, 0.04], (0.04, 0.25], (0.25, 0.5], (0.5, 1] as thresholds for the four-class RSA predictions (Buried, Partially Buried, Partially Accessible, Accessible), and [0, 0.25], (0.25, 1] as thresholds for the two-class RSA predictions (Buried, Accessible). We discarded the proteins for which DSSP could not produce an output, as well as those that had irregularities in their PDB formats. The final set contained 9,680 protein chains.

#### 2.4.4 Quality Measures

The measure of the performance of the trained ensemble of SS-RSA networks is the three-class accuracy (Q3) for SS and the four-class and two-class accuracy for RSA, which are calculated by dividing the number of correctly classified residues by the total number of residues in the dataset.

CMAP predictions for the CASP12 RR benchmark set were evaluated in accordance with the CASP criteria by calculating the accuracy of the top *L*/5 predicted long-range contacts, where *L* is the length of the protein, and the long-range contacts are contacts between residues with sequence separation distance over 23.

#### 2.4.5 Ensembling

Ensembling models usually yield a consensus model that performs better than any of the networks included in the ensemble [63]. Several networks both for CMAP and SS-RSA have been trained with different parameters (see “Results” section). Even though some models have worse performances on average, they are still saved. All the saved models that have been trained on the same set are used at testing time. The outputs from each model are ensembled to determine the final output. This is done by averaging all outputs from the softmax units and selecting the final class by picking the class with the highest average probability.

In the CMAP case, each model in the ensemble is used to make two predictions for each target using either HHblits or JackHMMER alignment. Although the CMAP network is trained only on HHblits alignments, using the JackHMMER alignments in the ensemble improved the overall accuracy of the predictor.

## 3 Results and discussion

### 3.1 Embeddings

Tensorboard in Tensorflow can be used to visualize the output of the embeddings layer to see how each residue type is mapped on the output space. In Fig. 2 we show the embeddings outputs for an early version of the SS-RSA network where each amino acid type is mapped on a 4D space. The 4D vectors are projected onto a 2D space by principal component analysis on Tensorboard. In the figure, the amino acids that are the closest (lowest cosine between the 4D vectors) to Lysine (Fig. 2-a) and Tryptophan (Fig. 2-b) are highlighted. The amino acids closest to Lysine (K) are Histidine (H) and Arginine (R), which makes sense, since they can all be positively charged. Similarly, the residues closest to the hydrophobic Tryptophan (W) are also hydrophobic, indicating that the embeddings can discriminate between different kinds of amino acids and map them onto a space that makes sense from a chemical point of view.

**Fig 2.**
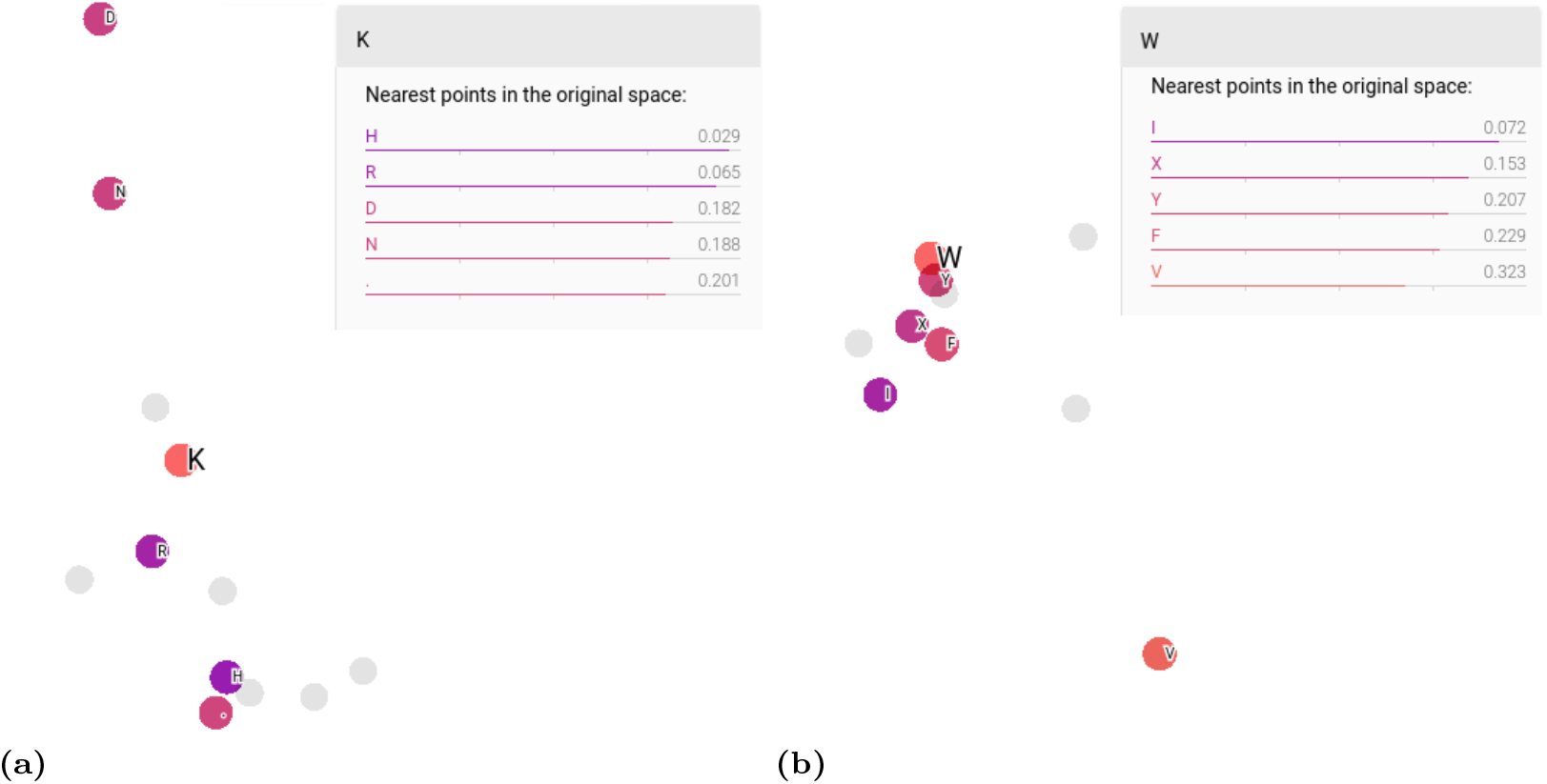
2D PCA of the space of the embedded vectors representing the single residues. The original space has a dimensionality of four. The residues that are closest (lower cosine between the 4D vectors) to (a) Lysine and (b) Tryptophan are colored (the closer the residue, the darker the hue).

### 3.2 SS and RSA predictions

We have used the five-fold cross-validation results to determine the testing accuracy for SS-RSA. To compare the rawMSA approach against a classic profile-based method, we trained a separate network by removing the bottom layers from the SS-RSA network (embedding and first 2D convolution/Pooling layer) and we trained it by using the PSSMs calculated from the HHblits alignments as inputs. We tested and trained the PSSM network in the same way we trained the other networks, both for SS and RSA. In Fig. 3 we compare the performance of the PSSM network against several rawMSA networks trained on different numbers of input sequences (100 to 1000 MSA sequences). The boxplot shows how the rawMSA networks with more input sequences perform generally better, with the rawMSA500 and rawMSA1000 networks performing slightly better than the classic PSSM network in predicting secondary structure, and all the rawMSA networks outperforming the PSSM network in predicting solvent accessibility.

**Fig 3.**
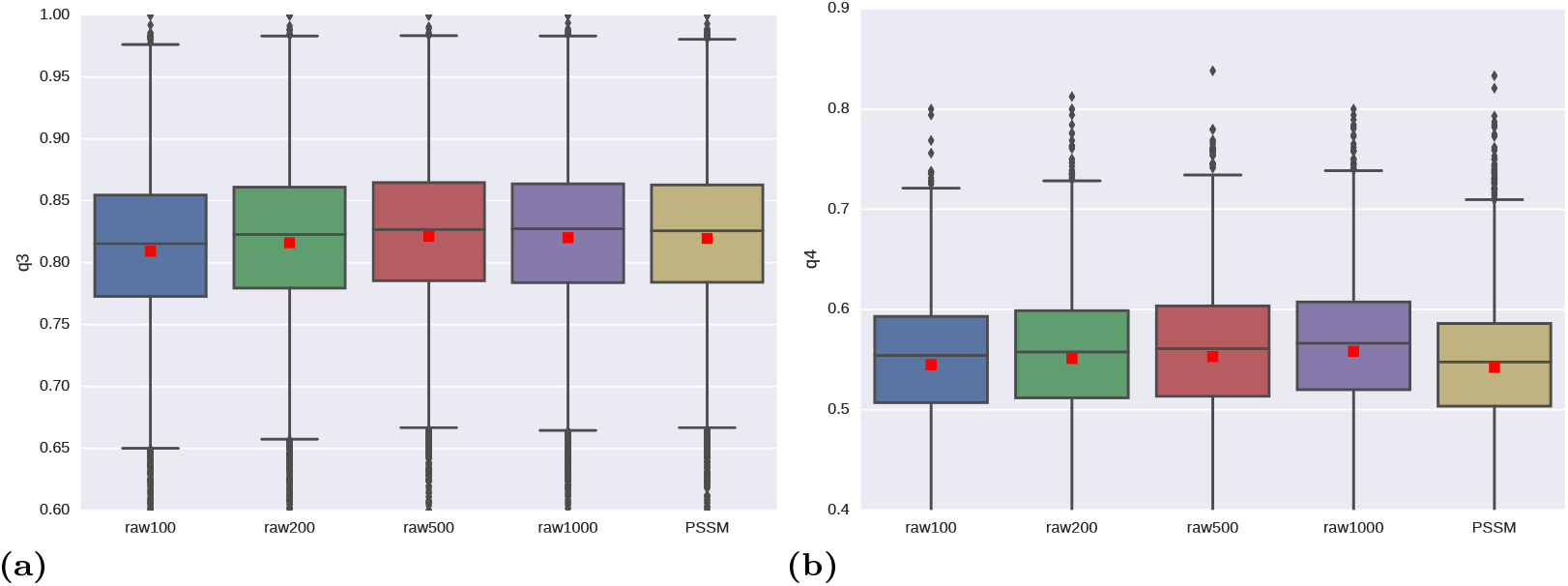
Boxplots showing the accuracy distribution (target-wise) for the predictions by a number of rawMSA networks against a classical PSSM network trained and tested on the same dataset. Here are shown the results both for secondary structure (a) and four-class solvent accessibility (b) predictions. Four different rawMSA networks are tested at variable MSA depths (number of top alignments considered from the HHblits MSAs as input to the network). Here, the top 100, 200, 500 or 1000 alignments from the MSAs are given as input to the SS-RSA network. The average accuracies are also shown (red squares).

The final SS-RSA network is an ensemble of six networks trained in five-fold cross-validation on 100 to 3000 input MSA sequences per target.

The results are shown in Table 1.

**Table 1.** Accuracy results for the SS-RSA network trained to predict secondary structure and solvent accessibility (2-class and 4-class).

| Predictor | Accuracy |
| --- | --- |
| rawMSA SS ensemble | 83.4 |
| rawMSA1000 SS | 81.8 |
| PSSM SS | 81.7 |
| rawMSA RSA ensemble (4-class) | 57.7 |
| rawMSA1000 RSA (4-class) | 56.1 |
| PSSM RSA (4-class) | 54.1 |
| rawMSA RSA ensemble (2-class) | 81.2 |

It is difficult to make a direct comparison of rawMSA against other predictors in literature because of inevitable differences in the datasets. One example of this comes from secondary structure prediction systems, which have recently been reported to predict at accuracies (Q3) of up to 84% [64], yet we have not been able to find a recent study where the reported accuracy is supported by a proper splitting of the training and testing sets (see “Avoiding homolog contamination” paragraph). Running local versions of existing software does not solve that problem since it is not clear exactly which sequences were used for training. Also, in many cases a final network is trained using all available sequence, which means that any test is bound to be contaminated by homologous information. However, given the very large size of our test set, the rigorousness of our experimental setup, and the fact that rawMSA outperforms our own PSSM-based method, we believe that rawMSA compares favorably against the state of the art.

### 3.3 CMAP predictions

The final CMAP network is an ensemble of 10 networks trained on 10 to 1000 input sequences and varying numbers of layers (10 to 24 convolutional layers). The CMAP predictions for each target have been sorted by the contact probability measure output by the ensemble, then the top *L*/5 long-range contacts have been evaluated against the native contact map. The final accuracy has been calculated as the average of the accuracies for all targets. In order to make a fair comparison against the other predictors, we have downloaded all of the predictions made in CASP12 and evaluated them with the same system. In Table 2 we compare the top *L*/5 long-range accuracy of rawMSA CMAP against the top 5 CASP12 predictors.

**Table 2.** Comparison of rawMSA against the top 5 contact prediction methods in CASP12. All predictions from CASP12 have been re-evaluated to ensure a fair comparison. The accuracy is calculated on the top L/5, long range contacts.

| Predictor | Domain Count | L/5 LR Accuracy |
| --- | --- | --- |
| **rawMSA CMAP** | **37** | **43.8** |
| RaptorX-Contact | 37 | 43.0 |
| iFold_1 | 36 | 42.3 |
| Deepfold-Contact | 37 | 38.6 |
| MetaPSICOV | 37 | 38.4 |
| MULTICOM-CLUSTER | 37 | 37.9 |

rawMSA outperforms the top predictors in CASP12 under the same testing conditions. This is especially impressive since it is the only top predictor not to use any kind of explicit coevolution-based features, or any other inputs other than the MSA. It is important to note that only up to 1000 input MSA sequences could be used in training and testing because of limits in the amount of GPU RAM available (*<*25GB). Since there is a correlation between the testing accuracy and the number of sequences used as input, it is reasonable to expect that rawMSA will benefit from training on GPUs with larger memory.

## 4 Conclusion

We have presented a new paradigm for the prediction of structural features of proteins called rawMSA, which involves using raw multiple sequence alignments (MSA) of proteins as input instead of compressing them into protein sequence profiles, as is common practice today. Furthermore, rawMSA does not need any other manually designed or otherwise hand-picked extra feature as input, but instead exploits the capability that deep networks have of automatically extracting any relevant feature from the raw data.

To convert MSAs, which could be described as categorical data, to a more machine-friendly format, rawMSA adopts embeddings, a technique from the field of Natural Language Processing to adaptively map discrete inputs from a dictionary of symbols into vectors in a continuous space.

To showcase our novel representation of the MSA, we developed a few different flavors of rawMSA to predict secondary structure, relative solvent accessibility and inter-residue contact maps. All these networks use the same and only kind of input, i.e. the MSA. After rigorous testing, we show how rawMSA SS-RSA sets a new state of the art for these kinds of predictions, and rawMSA CMAP performs on par with the top predictors from the CASP12 experiment even though relatively few alignments (up to 1,000 per protein in this experiment) could be used as input, due to limitations in the amount of available GPU RAM.

We believe that rawMSA represents a very promising and powerful approach to protein structure prediction, and with the ever-increasing amounts of computational power and RAM available in the next-generation GPU architectures, it could very well replace older methods based on protein sequence profiles in the coming years.

## Acknowledgement

This work was supported by Swedish Research Council grants 2012-5270, 2016-05369, The Swedish e-Science Research Center, and the Foundation Blanceflor Boncompagni Ludovisi, née Bildt. Computations were performed on resources provided by the Swedish National Infrastructure for Computing (SNIC) at the National Supercomputer Centre (NSC) in Linköping and at the High Performance Computing Center North (HPC2N) in UmeÅ. We would also like to thank Hops (www.hops.io), for providing some extra GPU resources, while others were graciously provided by NVIDIA Corporation through the GPU Grant Program. We also thank Isak Johansson-Åkhe for helpful discussions.

